# Stochastic differential equation modelling of cancer cell migration and tissue invasion

**DOI:** 10.1101/2022.11.14.516390

**Authors:** Dimitrios Katsaounis, Mark A.J. Chaplain, Nikolaos Sfakianakis

**Author notes:** Contributing authors.

## Abstract

Invasion of the surrounding tissue is a key aspect of cancer growth and spread involving a coordinated effort between cell migration and matrix degradation, and has been the subject of mathematical modelling for almost 30 years. In this current paper we address a long-standing question in the field of cancer cell migration modelling. Namely, identify the migratory pattern and spread of individual cancer cells, or small clusters of cancer cells, when the macroscopic evolution of the cancer cell colony is dictated by a specific partial differential equation (PDE).

We show that the usual heuristic understanding of the diffusion and advection terms of the PDE being one-to-one responsible for the random and biased motion of the solitary cancer cells, respectively, is not precise. On the contrary, we show that the drift term of the correct stochastic differential equation (SDE) scheme that dictates the individual cancer cell migration, should account also for the divergence of the diffusion of the PDE. We support our claims with a number of numerical experiments and computational simulations.

## 1 Introduction

Cancer invasion is a complex process involving numerous interactions between the cancer cells and the extracellular matrix (ECM) (cf. the tumour microenvironment) facilitated by matrix degrading enzymes. Along with active cell migration (both individual and collective) and increased/excessive proliferation, these processes enable the local spread of cancer cells into the surrounding tissue. Any encounter with blood or lymphatic vessels (cf. tumour-induced angiogenesis, lymph-angiogenesis) in the tumour microenvironment initiates and facilitates the spread of the cancer to secondary locations in the host, i.e., metastasis or metastatic spread. A comprehensive historical overview of the biology of metastastic spread can be found in the article by Talmadge and Fidler [1], while an overview of the core aspects of invasion can be found in the articles of Hanahan and Weinberg [2], [3] and the review article of Friedl and Wolf [4]. From a mathematical modelling perspective, cancer invasion has been a topic of interest for almost 30 years with a range of approaches and techniques being used, and an overview can be found in the recent review paper by Sfakianakis and Chaplain [5]. Broadly speaking, two different approaches have been used to model cancer invasion-continuum approaches (i.e. using differential equations with cancer cell density as one of the dependent variables) and individual-based or agent-based approaches (i.e. focusing on the movement of individual cells). Some have also adopted a so-called hybrid approach e.g. [6], deriving a discrete model governing the migration of individual cancer cells from the discretization of an associated system of PDEs.

It is not our intention here to discuss in detail the previous modelling work in the area. Rather the aim of this paper is to investigate mathematically the connection between the stochastic differential and the partial differential equations (SDEs and PDEs respectively) that are typically used to describe the migration of living cells. The precise interplay between the SDE and PDE approach is not yet clear and has been in the research focus the last years. The difference between these two approaches is significant and it lies primarily in the immediate focus of the mathematical model. Namely, whereas the SDE approach focuses primarily on the migratory behaviour of the individual cells, the PDE approach describes the macroscopic behavioural pattern of a large collective of cells. As the behavioural pattern, in real life biology, of large cell collectives is related to the migration of individual cells so should the two mathematical approaches be connected.

By identifying the interplay between the two mathematical approaches, we shed light in the complexity of multiscale modelling and simulations. This has direct implications in the biological understanding of solitary cancer cell migration and the development/growth of tumours. Such detailed knowledge of how far individual cancer cells can penetrate into the local tissue is very important from a surgical point of view and can help to minimise the amount of resection required, a point initially raised and investigated in the work of Anderson et al. [6].

Our aim, hence, in this work is to investigate the interplay between the SDE and PDE modeling approaches of cancer invasion and growth. Namely, we exhibit that the terms of the numerical scheme solving the underlying SDE are not in a one-to-one correspondence to the terms of the PDE. In more detail, we show that a particular “correction” in the drift terms of the numerical scheme of the SDE that improves the approximation qualities of the schemes when compared with the numerical solution of the PDE.

These ideas are studied in the remainder paper in the following way. In Section 2 we provide some background for the motivation to this paper stating the general forms of PDE and SDE to be considered, while in Section 3 we derive the SDE schemes in some detail. In Section 4 we undertake numerical experiments and compare results from computational simulations of the underlying cell migration PDE model with simulations of two different SDE schemes. Finally in Section 5 concluding remarks are made.

## 2 Motivation

We are motivated in this work by typical continuum cancer invasion models (e.g. [7], [8], [9]) and consider the following general Advection-Diffusion PDE

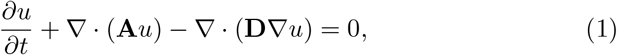

where, according to the usual practice, *u* : [0, *T*] × Ω → ℝ represents the spacetime dependent density of cancer cells, and where Ω ⊂ ℝ^*d*^, with *d* = 2, 3, is a Lipschitz domain. We assume throughout this work that both the advection and diffusion coefficients **A** and **D** are non-constant in the sense that they depend on **x** ∈ ℝ^*d*^.

It is biologically understood that the macroscopic patterns of a large collective of cells is related to the migration of individual cells. Hence, our aim is to understand the connection between the continuous model (1), and models that capture the migration of solitary cancer cells. Following the ideas developed in the seminal works by Einstein [10] on the investigation of Brownian motion, as well as by Stratonovich, Ito and Kitanidis [11–13], the motion of the solitary cancer cells is usually described via SDEs that track the position of the cells. These SDEs typically take the form

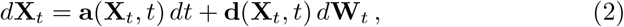

with *t* ≥ 0. Here **X**_*t*_ ∈ ℝ^*d*^ is a stochastic process that represents the position of the solitary cancer cells, and where **a**(**X**_*t*_, *t*) and **d**(**X**_*t*_, *t*) are the drift and the diffusion coefficients respectively. Here **W**_*t*_ represents a *d*-dimensional *Wiener process*.

The critical question that arises in this work is the following: how do the advection and diffusion terms **A, D** of the deterministic PDE (1) relate to the drift and diffusion terms **a, d** of the SDE (2)?

By identifying these relations we shed light in the complexity of multiscale modelling and simulations in general, and in the migration of solitary cancer cells and macroscopic tumour growth within a living organism in particular. Succeeding, hence, in identifying such relations will allow well established— and phenomenologically verifiable—macroscopic models to be used in order to extract, at the smaller scale, the migratory pattern of solitary cells or small clusters of cancer cells. Vice-versa, these relations will provide an additional validation to the use of deterministic models or even encourage ideas for a hybrid method using both deterministic and stochastic models.

We proceed in this effort under the assumption that *u* represents a single solitary cancer cell or a small cluster of cancer cells. We denote by **x**_*t*_ ∈ ℝ^*d*^ the numerical approximation of the solution stochastic process **X**_*t*_ ∈ ℝ^*d*^that represents the position of the cell’s centroid at time *t*. Following [11–15] we exhibit in Section 3 that the *less* intuitive numerical scheme:

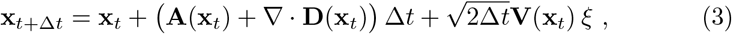

where ***ξ*** is a vector of *d* independent and normally distributed random variables of mean 0 and variance 1, and where **V** given by

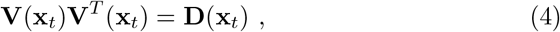

provides a better approximation to the PDE (1) than the *more* intuitive numerical scheme:

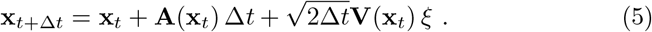

Both schemes account, in the same way, for the “square root” of the diffusion **D** in their corresponding diffusion/noise terms. As we will see in Section 3 this is derived in a very natural way.

The SDE (2) is of an Ito-type and, accordingly, the schemes (3) and (5) are derived using the Euler-Maruyama approximation. The less intuitive scheme (3) should not be mistaken for the equivalent scheme of a Stratonovich-type SDE emerging from (2), cf. [15]; it should rather be understood as an Ito-type scheme that is different from (5).

Structurally, the difference of the two schemes (3) and (5) lies in their corresponding drift terms. In particular, in the “corrected” scheme (3) the drift term accounts for both the advection, **A**, and the diffusion, **D**, coefficients of the PDE (1). In the scheme (5) the drift term accounts only for the advection term **A**. This final remark, the one-to-one correspondence between the terms of the scheme (5) and the equation (2), justifies the characterisation of the scheme (5) as “more intuitive”, cf. [15, 16].

In Section 3 we derive the scheme (3) from the PDE (1) and in Section 4 we compare it numerically with the scheme (5) and the PDE (1).

We, hence, conclude that there is not a one-to-one relation between the advection term of the PDE and the drift term of the SDE—unless the divergence of the diffusion **D** vanishes; this is clearly the case when the diffusion **D** does not depend on the spatial variable **x**.

## 3 Derivation of the SDE Schemes

We make the fundamental modeling assumption that a cell can be viewed as a sufficiently small cell-cluster that satisfies the PDE (1). We furthermore assume that, without loss of generality, such a cell-cluster has unit mass i.e.

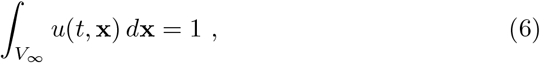

where *u*(*t*, **x**) is the density and *V*_*∞*_ the volume of the cell-cluster.

To capture the behaviour of the cell-cluster, and in particular, the position of its centroid and spread, we use the method of moments. Accordingly, the first and second moments of *u* read respectively

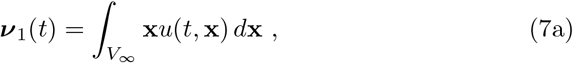

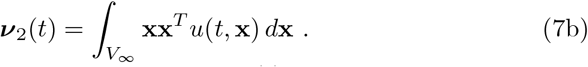

Note that ***ν***_1_ (*t*) is a *d*-dimensional vector and ***ν***_2_ (*t*) a *d* ×*d*-dimensional matrix, and they are both understood via their physical rather than probabilistic interpretation. We also consider the second moment about the mean

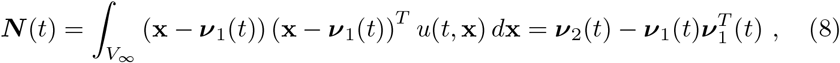

where ***N*** (*t*) is *d × d*-dimensional matrix. The first moment ***ν***_1_ in (7a) can be interpreted as the position of the cell-cluster centroid, and, accordingly, it’s (time) rate of change as the velocity of the cell-cluster. On the other hand, the rate of change of the second moment ***ν***_2_ in (7b) represents the spreading of the cell-cluster. These two remarks together allow to (heuristically) identify the drift and diffusion coefficients, **a** and **d**, of the SDE (2) as 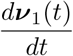 and 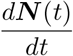 respectively.

As a direct consequence of that, we will construct a numerical scheme in line with the classical Euler-Maruyama approximation, cf. [14], as follows

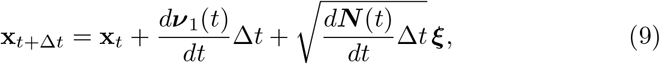

where the square root in the above, should be understood in the usual matrix notation, see e.g. (4).

To this end, we impose the following boundary conditions that indicate an exponential decay of *u*(*t*, **x**) as |**x**| → *∞*.

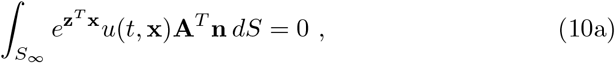

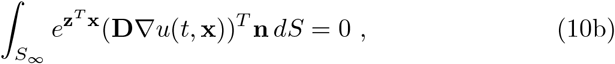

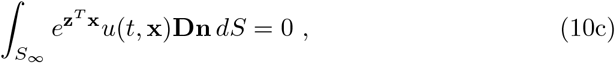

where *S*_∞_ represents the surface of the cell-cluster *V*_∞_ and **z** a *d*-dimensional vector. Through the *i*-th derivative of (10a)-(10c) with respect to **z** and setting **z** = 0 one can retrieve the boundary conditions for the *i*-th moment. In our case we only need to differentiate twice since we only need the first two moments.

With all these in mind, we proceed by calculating through (7a) and (8) the time derivatives of ***ν***_1_ (*t*) and ***N*** (*t*). For ***ν***_1_ we multiply (1) by *x*_*i*_, for *i* = 1, …, *n*, integrate over the cell-cluster volume *V*_∞_, and obtain

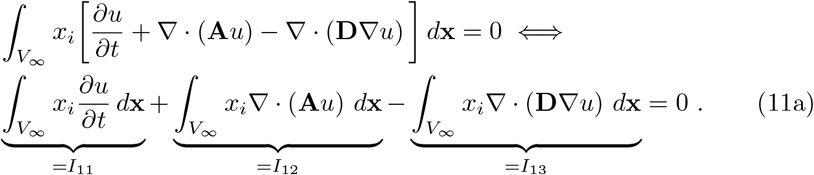

We work on each of the terms *I*_11_, *I*_12_, *I*_13_ on the left hand side separately; and obtain for *I*_11_

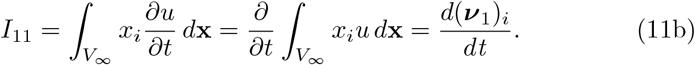

For the second term *I*_12_, after invoking the Divergence Theorem and the boundary conditions (10a)-(10c), we obtain

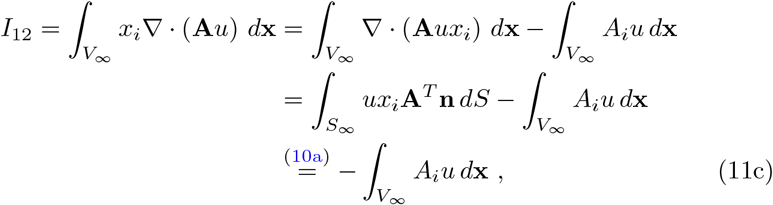

where *A*_*i*_ is the *i*-th element of the vector **A**. Similarly, the third term *I*_13_ recasts into

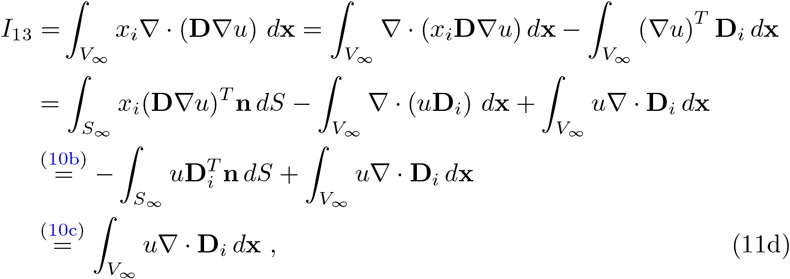

where **D**_*i*_ is the *i*-th column of the diffusion matrix **D**. So, by combining (11a) with (11b)-(11d) we obtain the following representation for the rate of change of the *i*-th (vector) component of the first moment:

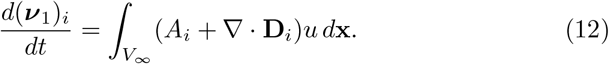

In a similar way, we identify the relation satisfied by the *ij*^th^ element of the second moment ***ν***_2_, which allows us to compute the rate of change of ***N*** (*t*) in (8):

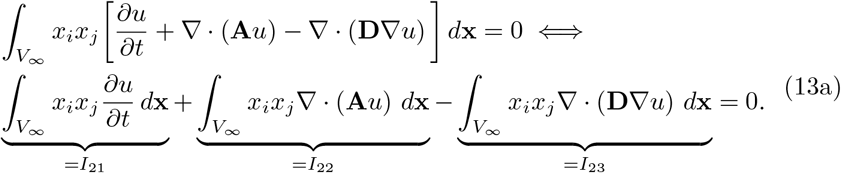

As with (11a), we employ the Divergence Theorem and the boundary condition (10a)-(10c) and calculate these three integrals one-by-one. The first one, *I*_21_, reads

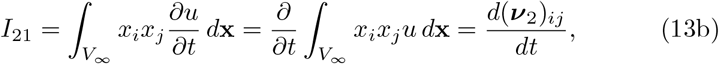

the second integral, *I*_21_, recasts into

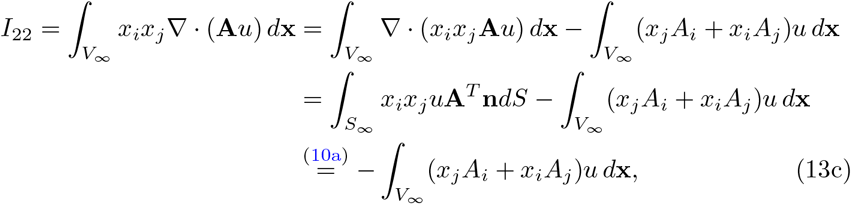

and for the third term, *I*_21_, it holds

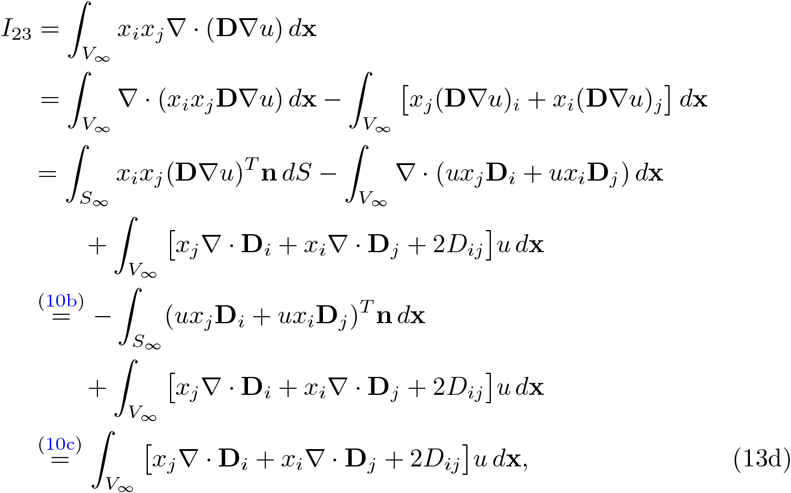

where *D*_*ij*_ is the element in the *i*-th row and *j*-th column of the matrix **D**. By combining (13a) with (13b)-(13d), we obtain the following relation for *ij*-th (matrix) element of the second moment

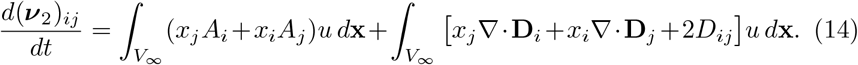

Summing up (12) and (14), we deduce the following set of equations in a vector form for any given advection and diffusion terms **A** and **D** in (1)

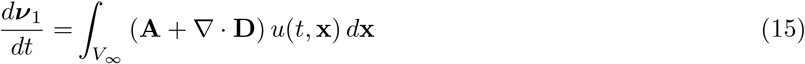

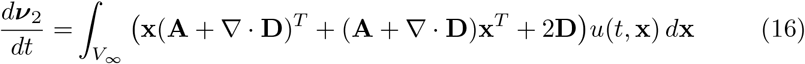

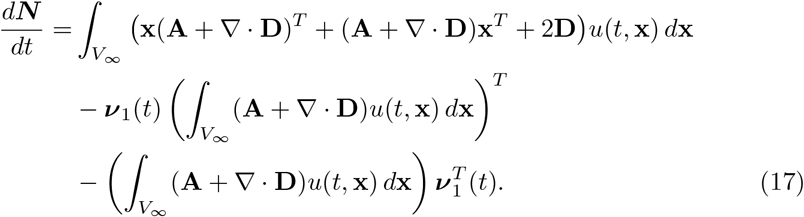

We retrieve (17) by taking the time derivative of (8) and substituting equations (15) and (16).

Assuming that the mass of the cell-cluster is concentrated, at time *t*, in a single point **x**_*t*_, we can represent the density of the cell-cluster as

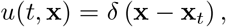

where *δ* is the Dirac function centered at **x**_*t*_. Then the velocity of the cell-cluster in (15), is given as the sum of the advection **A** and the divergence of the diffusion matrix **D** at **x**_*t*_ :

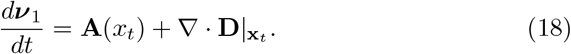

Accordingly, the rate of change of second moment about the mean in (17) recasts into

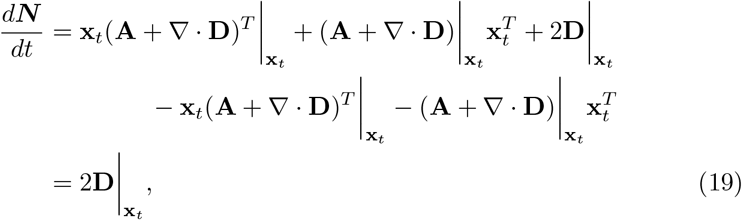

which clearly indicates that the rate of spreading of the cell-cluster is given by the diffusion matrix at the point **x**_*t*_ .

We close this section by substituting (18) and (19) into (9) to obtain the corrected numerical scheme

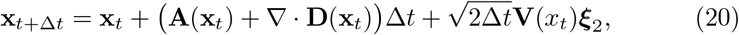

where **V**(**x**_*t*_)**V**^*T*^ (**x**_*t*_) = **D**(**x**_*t*_) and where ***ξ***_2_ is a vector of *d* independent and normally distributed random variables with 0 mean and variance 1.

The scheme (20) that we have just derived, is different from the one without the correction of the drift term, i.e. (5), which we repeat here for completion:

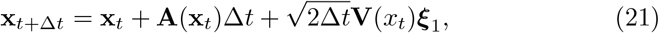

where, as before, ***ξ***_1_ is a vector of independent and normally distributed random variables with mean 0 and variance 1.

In the next section we numerically investigate the two schemes (21) and (20) and provide evidence of their differences and their fitting with the corresponding/underlying PDE (1).

## 4 Numerical Experiments

We have seen in the previous section that the drift term of the SDE scheme (20) incorporates a diffusion-based contribution/correction that is not found in the more intuitive scheme (21). This is a significant difference between the two schemes and is central in our numerical investigations. Namely our aim in this section is to numerically investigate the impact that this correction has on the simulations of these two schemes. To this end, we consider three specific numerical settings that highlight the difference of the two schemes, and compare their predictions with each other and with the corresponding/underlying PDE (1).

In more detail, in Experiment 1, we investigate the behaviour of the two schemes, (21) and (20), on a particular application where the corresponding noise terms ***ξ***_1_, ***ξ***_2_ are the same. This allows to csompare the two schemes as a result of their differences on the drift terms alone.

In Experiment 2, we consider the same computational setting as in Experiment 1, with the only difference being that the two SDE schemes (21), (20) are augmented with different noise terms ***ξ***_1_, ***ξ***_2_. With a large number of scheme realisations we extract information on the full spectrum of differences between the two schemes.

In Experiment 3 we consider a more generic, and common in the cancer invasion modelling literature, experimental setting where the advection and diffusion terms **A, D** of (1) depend on the spatial variable **x** ∈ ℝ^*d*^ through their dependence on the non-uniform tumour microenvironment *v* : ℝ^*d*^ → ℝ. For both SDE schemes (21) and (20), we perform a large number of realisations and compare them with the solution of the underlying PDE (1). We accordingly conclude that the corrected SDE scheme (20) provides a much better approximation to the PDE (1) than the more intuitive SDE scheme (21).

For the numerical solution of the PDE (1) we use a numerical method that was previously developed in [17, 18] and which we briefly discuss in the Appendix A. All algorithm implementations, simulations, and visualisations were conducted in MATLAB [19].

### Experiment 1

In the first experiment, we consider a modelling setting where the advection and diffusion terms, **A** and **D**, of the PDE (1) have an explicit dependence on the space variable **x**, namely:

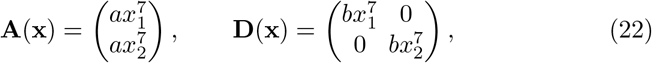

where **x** = (*x*_1_, *x*_2_) ∈ ℝ^2^ and *a, b* ∈ R are constants. Accordingly, the PDE (1) reads as

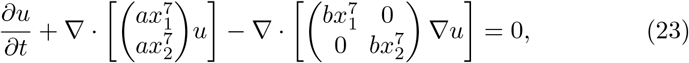

where *t* ∈ [0, *T*]. Similarly, the SDE schemes (21) and (20) are re-formulated, for

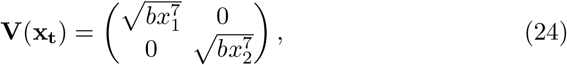

as follows:

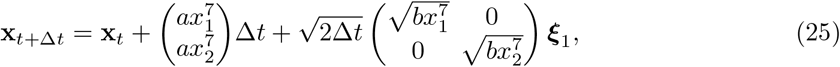

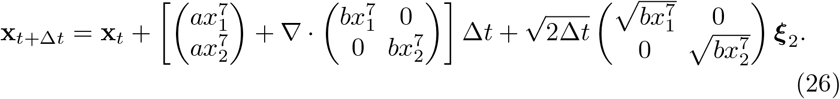

These two SDE schemes are augmented with the same noise terms ***ξ***_1_, ***ξ***_2_ to allow for a direct, one-to-one, comparison of their realisations.

The parameters used in this numerical experiment can be found in Table 1 and the simulation results are presented in Figure 1. In more detail, we perform 100 realisations of the two schemes (25), (26)—each pair of experiments with the same noise—starting from the same initial position (*x*_1_, *x*_2_) = (1, 1) and running over the time *t* ∈ [0, 1]. These results show that the realisations of the two schemes appear to be—in a one-to-one conformation—slightly shifted and parallel to each other. More specifically, as can be seen, the corrected scheme (26) introduces an additional displacement of the cell migration track/sample paths directed towards larger values of *x*_1_ and *x*_2_. This is due to the nature and structure of the drift term as well as the diffusion-based correction introduced in (26) and the positivity of the parameter *b*, cf. Table 1.

**Table 1.** Parameters used in **Experiments** 1 and 2.

| - | $t_0$ | T | N | $a$ | $b$ | $\mathbf{x}_0$ | realisations |
| --- | --- | --- | --- | --- | --- | --- | --- |
| Experiment **1** | 0 | 1 | 1000 | 0.01 | 0.001 | $(1, 1)^T$ | $10^2$ |
| Experiment **2** | 0 | 1 | 1000 | 0.01 | 0.001 | $(1, 1)^T$ | $10^5$ |

**Fig. 1.**
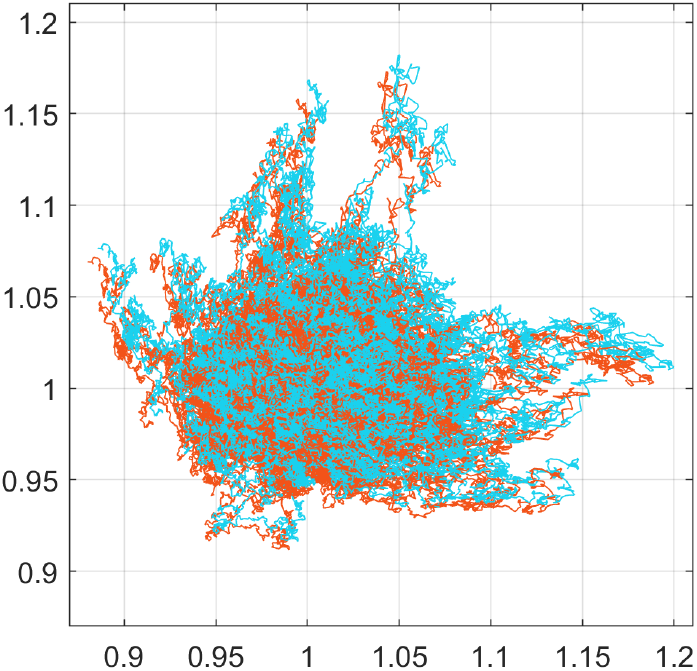
**Exp. 1**. Showing here the cell migration tracks/sample paths of 100 realisations of the SDE schemes (25), in orange, and (26), in light blue, with the same random noise. The initial position of the cells is set at (1,1) for all realisations and the total travelling time is *T* = 1. Note that the sample paths of the two schemes are parallel shifts of one another. The shift between the two schemes is due to the adaptation of the drift term of the scheme (26).

### Experiment 2

The intuition we have gained from Experiment 1, namely the way the adaptation of the drift term in the corrected scheme (26) affects the distance and direction of the cell migration, can be investigated further by considering independent noise terms for the two schemes. Hence, in the current experiment, we choose independent noise terms ***ξ***_1_, ***ξ***_2_ for the two SDE schemes (25) and (26) and perform 10^5^ new realisations with each, all start from the initial point (*x*_1_, *x*_2_) = (1, 1) and running over time *t* ∈ [0, 1]. We otherwise consider the same setting and parameters as in Experiment 1; the parameters for this experiment can be found in in Table 1.

The simulation results are presented in Figures 2 and 3. In Figure 2, in particular, we see that cells migrate further away from the origin and in a more biased fashion when they follow the corrected scheme (26) rather than the scheme (25). This is clearly the result of the additional bias introduced in the drift term of the corrected scheme (26). The qualitative difference between the two schemes can be further seen in Figure 3 where we present the final positions of the cells at time *T* of both schemes, along with their corresponding convex hulls.

**Fig. 2.**
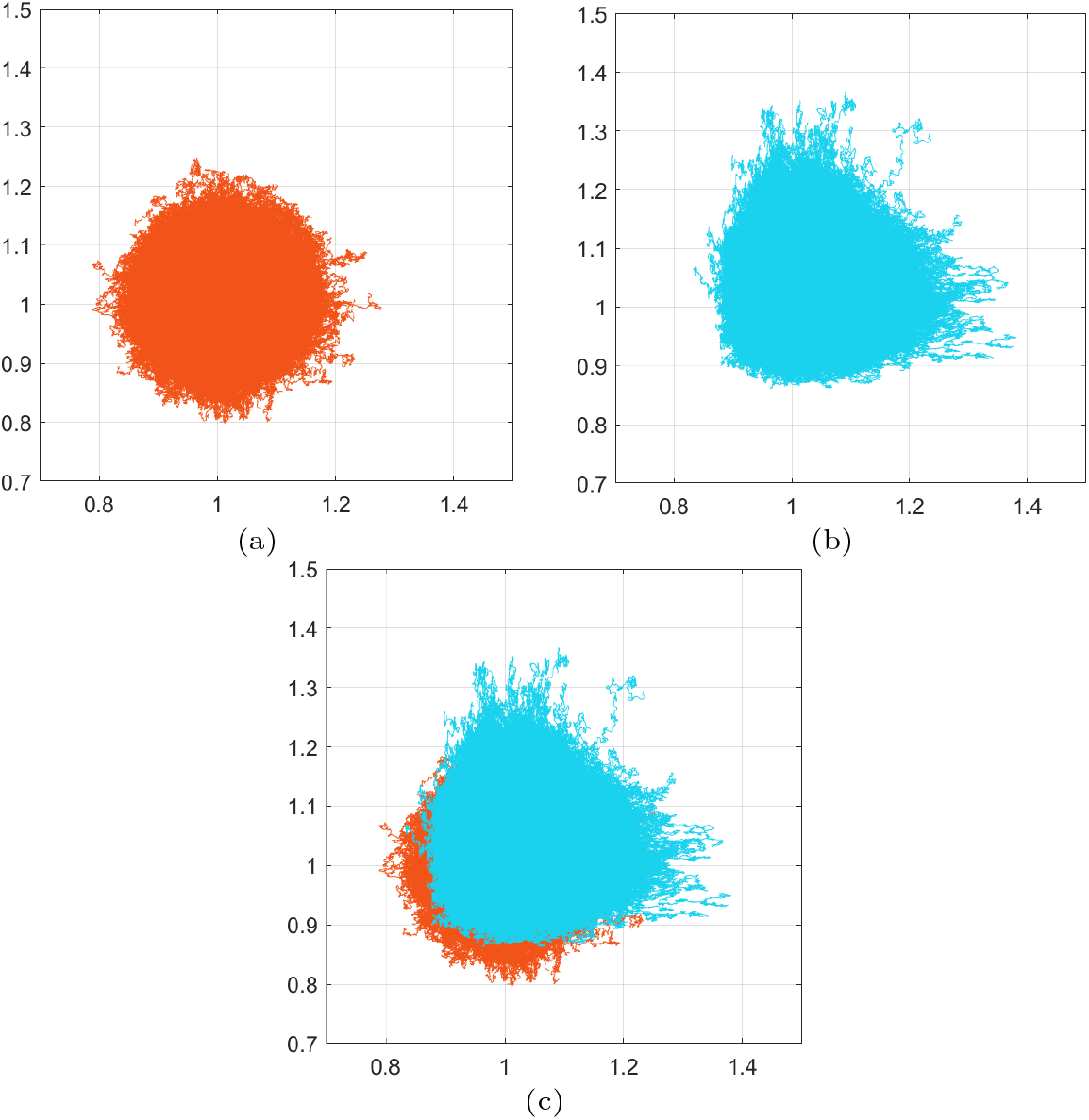
**Exp. 2**. Comparison of the migration tracks traversed the cells following the SDE schemes (25) and (26). For both schemes, we considered initial position at (1, 1) and a total travelling time *T* = 1. (a) & (b) The full migration tracks of 10^5^ cells following the SDE scheme (25) and (26) respectively. (c) Superimposing the migration patterns of (a) and (b) clearly reveals that the corrected scheme (26) introduces additional migratory bias.

**Fig. 3.**
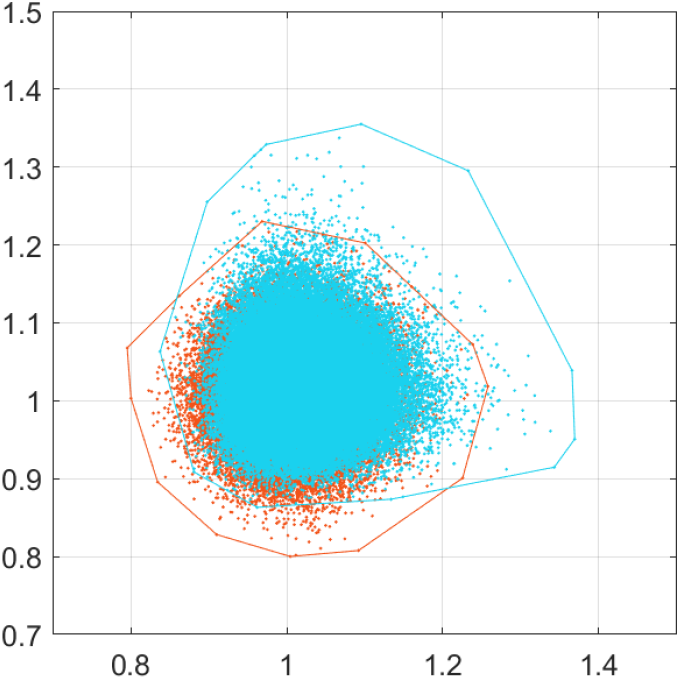
**Exp. 2**. Showing here the final positions of the cells migration after 10^5^ realisations of the SDE schemes (25) and (26)—in orange and light blue respectively—as shown in Figure 2. All realisations start from (1, 1) and run for time *t* ∈ [0, 1]. The corresponding convex hulls of these final positions are also shown. Note that the adaptation in the drift term of the corrected scheme (26), as opposed to the scheme (25), induces additional migration of the cells and spread of their final positions.

To measure the quantitative difference of the two schemes (25) and (26), we first measure the average distance traversed by the cells from their initial position until the final time *T*. In more detail, we perform *K* realisations of the schemes and calculate, for each one, the distance between the initial and the final position. We then calculate the average distance the cells have traversed by the formula

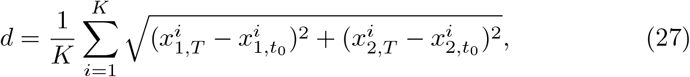

where 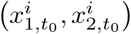 and 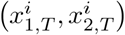 represent the initial and final positions of the cells in the realisation *i* = 1, …, *K*. We apply the above formula for the two schemes (25) and (26), after performing *K* = 10^6^ realisations of each, calculate their respective average distances *d*_1_ and *d*_2_ respectively, and evaluate the signed relative difference between the two to obtain

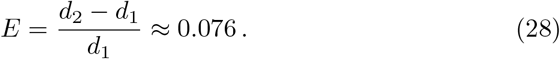

To further quantify the difference between the two schemes, we perform a uniform binning approach of the final positions of the cancer cells as described in [20]. Namely, we consider a partition of the *x*_1_ -axis into non-overlapping bins of fixed size *r*, and assign to each bin the quantity:

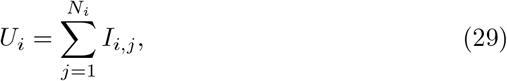

where *N*_*i*_ is the total number of positions (*x*_1_, *x*_2_) in the *i*-th bin and *I*_*i,j*_ is the *j*-th element of the set *I*_*i*_ = {*x*_2_ : for positions (*x*_1_, *x*_2_) in the *i*-th bin}.

The quantification was made by computing the *J*_2_ criterion which is defined as follows

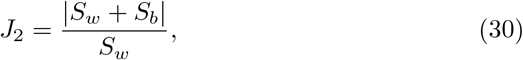

where

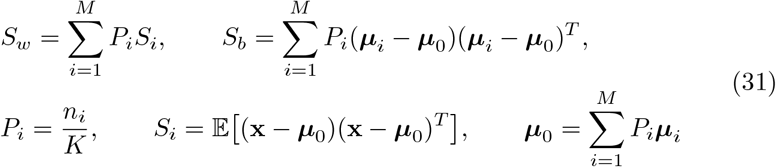

where *n*_*i*_ is the number of positions in the *i*-th bin, *K* is the number of realisations, and ***μ***_*i*_ is the mean value of *i*-th bin. After computing the *J*_2_ values of both schemes (25) and (26), we compute the signed relative error for different values *r* of the size of the bins

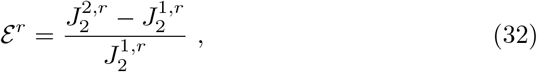

where 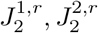, are the values of the *J*_2_ -criterion for schemes (25) and (26) respectively. Larger values of *J*_2_ correspond to better separated data. For the choice of *r* = 0.01 + *kh*, where *h* = 0.001, for *k* = 1, …, 60, we observe that for smaller sizes of the bins we get larger values of *J*_2_ and that 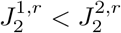 for all the values of *r*. The average value of the *k*–different values of *ε*^*6*^ is

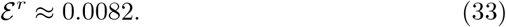

This result provides a second confirmation that (26) introduces additional migratory bias to the one side of the plane.

### Experiment 3

In this experiment we consider a PDE that is more common in the field of cancer invasion than the PDE in equation (23). Namely, we consider here a model where the growth of the tumour depends on the extracellular environment. This could represent, e.g. an extracellular chemical signal, the density of the extracellular matrix, or a completely different extracellular biochemical queue. Still, for the sake of simplicity of presentation, we do not make any particular biological assumptions on the nature of the extracellular environment and rather refer to it simply as “environment”.

Furthermore, we assume that this environment is non-uniform in space, does not change in time, and influences the growth of tumour in a very specific fashion. These assumptions are incorporated in the following model:

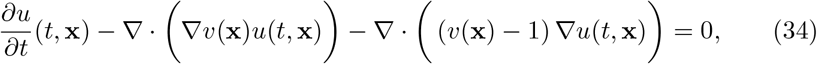

where *t* ≥ 0, **x** = (*x*_1_, *x*_2_) ∈ Ω = [−12, 12]^2^, and where *u* : [0, ∞) Ω → ℝ represents the density of the cancer cells. As previously mentioned, we do not investigate the biomedical realism of this model, nor do we interpret its findings under this light.

The PDE (34) is augmented with the initial condition

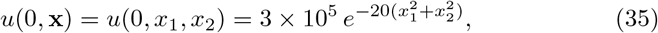

for **x** ∈ Ω as shown in Figure 4, and zero-Neumann boundary conditions over *∂*Ω. Furthermore, the (fixed) extracellular environment *v* : Ω → ℝ, shown in Figure 4, is given by

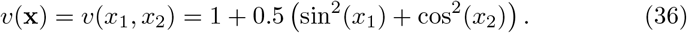

**Fig. 4.**
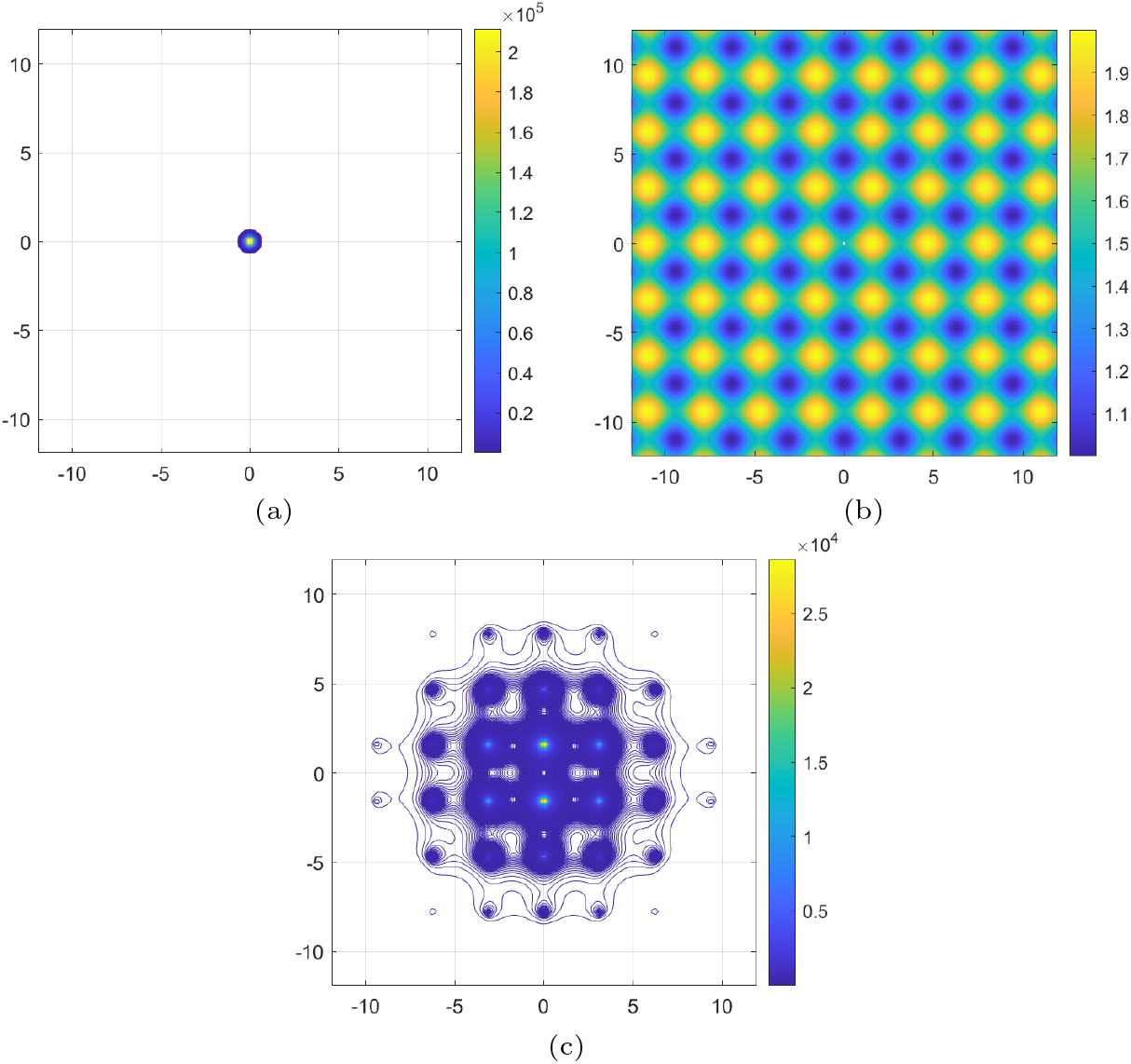
**Exp. 3**. Numerical solution of the PDE (34). (a) Showing here the isolines of the concentration *u* at time *t* = 0; they serve as initial conditions for the PDE (34). (b) The structure of the (fixed) extracellular environment *v* that participates in the advection and diffusion terms of (34); the formula of *v* is given in (36). (c) Isolines of the solution *u* of the PDE (34) at the final time *t* = 10; they reveal a higher concentration of the cancer cells in the “valleys” of the (fixed) extracellular environment *v* shown in (b).

Note that, for *v* given in (36), the diffusivity *v*(**x**) – 1 of (34) is non-negative for all **x** ∈ Ω.

We plot in Figure 4 the isolines of the solution of the PDE (34) at the final time *t* = 10. These illustrate clearly a significantly higher concentration of the cancer cells in the “valleys” of the extracellular environment *v*.

Based on the advection and diffusion terms of the PDE (34), we rewrite the SDE schemes (21) and (20) as follows:

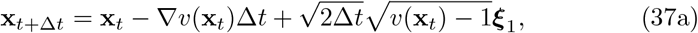

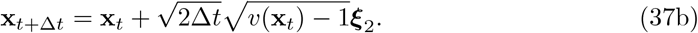

Note that the corrected SDE scheme (37b), which corresponds to the corrected scheme (20), lacks a drift term; this is a result of the particular structure of the advection and diffusion terms of the PDE (34) and the way they are combined in the drift term of the corrected scheme (20).

The simulations that we perform for Experiment 3 are shown in Figure 5 and are similar to the ones for Experiment 2 (cf. Figures 2 and 3). We note that the cell tracks of the more intuitive scheme (37a) spread out from the origin in a lesser extend than the corrected scheme (37b). This is due to the presence of the drift term in the scheme (37a) which leads almost all cancer cells to high densities of the extracellular environment *v*. On the other hand, the final positions of the sample paths given by (37b) concentrate less in the “valleys” of the extracellular environment.

**Fig. 5.**
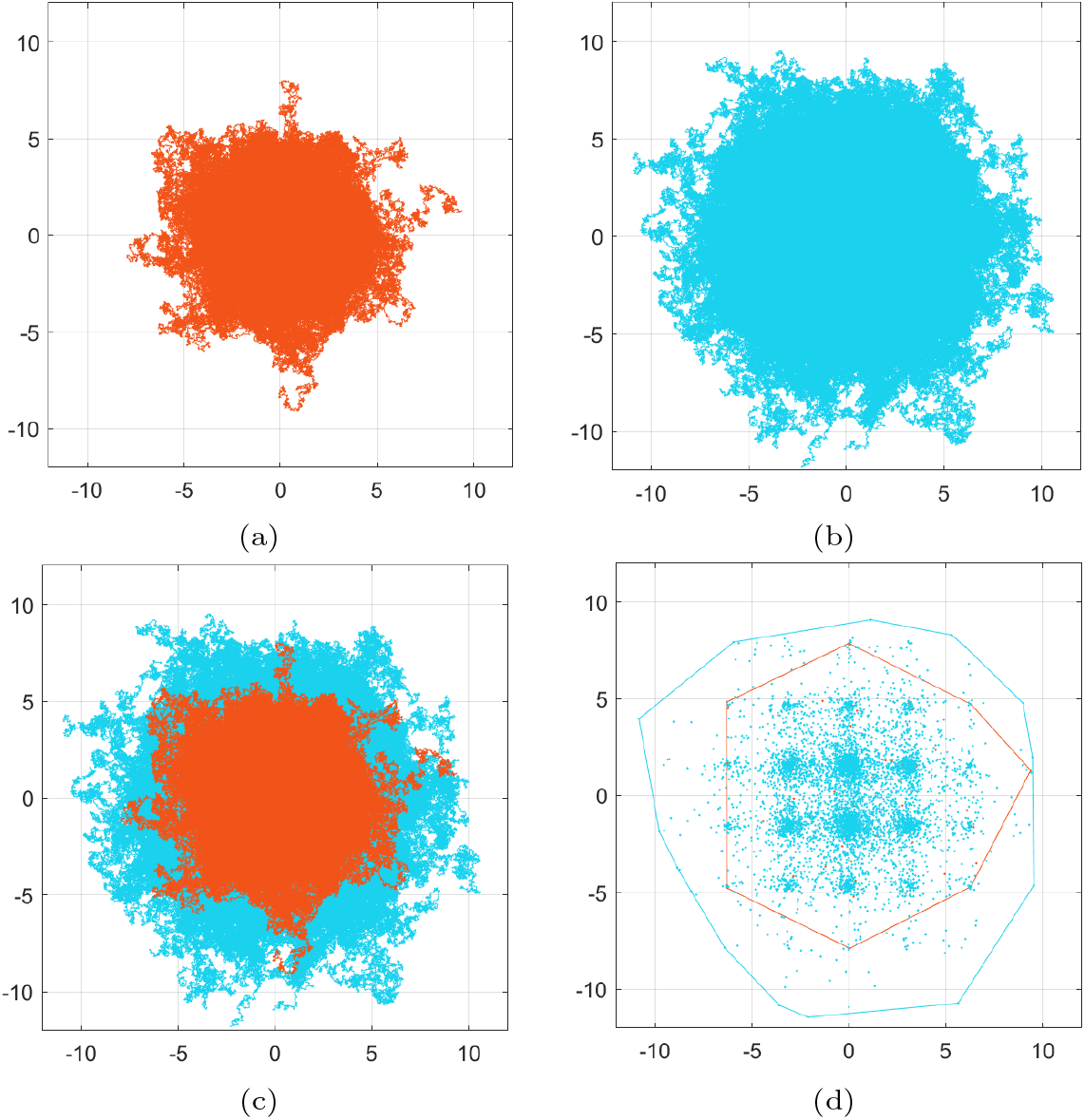
**Exp. 3**. Simulations and comparison of 10^4^ realisations of SDE schemes (37a) and (37b). (a) & (b) Full set of tracks/sample paths for the more intuitive scheme (37a) and the corrected scheme (37b) respectively. It can be clearly seen that the spread in (37b) is much wider than in (37a); this is justified by the adaptation introduced in the drift term of the corrected scheme (26). This remark is confirmed by superimposing the cell tracks of (a) in (b) in (c). (d) This shows the final positions of the 10^4^ realisations of (37a) and (37b), in orange and light blue respectively, along with the convex hulls of the corresponding points. It is clearly seen that the cells concentrate in the “valleys” of the environment *v*, cf. with the solution of the PDE (34) in Figure 4, and that the final positions of (37b) (shown in light blue) spread more than those of (37a). The final positions of (37a) are not visible as they are overlapped by the final positions of (37b).

In Figure 6 we present a direct qualitative comparison between the numerical solution of the PDE (34) and the SDE schemes (37a) and (37b). What this figure shows is that, at the final time *t* = 10, the positions of the cells of 10^4^ realisations of the SDE scheme (37a) are much more concentrated than the corresponding solution of the PDE (34). On the other hand, the distribution at the same final time *t* = 10, of 10^4^ realisations of the corrected scheme (37b) is more spread out and much closer to the solution of the PDE (34). In effect, these simulation results indicate that the corrected scheme (37b) is a better approximation to the numerical solution of the PDE (34) than the scheme (37a).

**Fig. 6.**
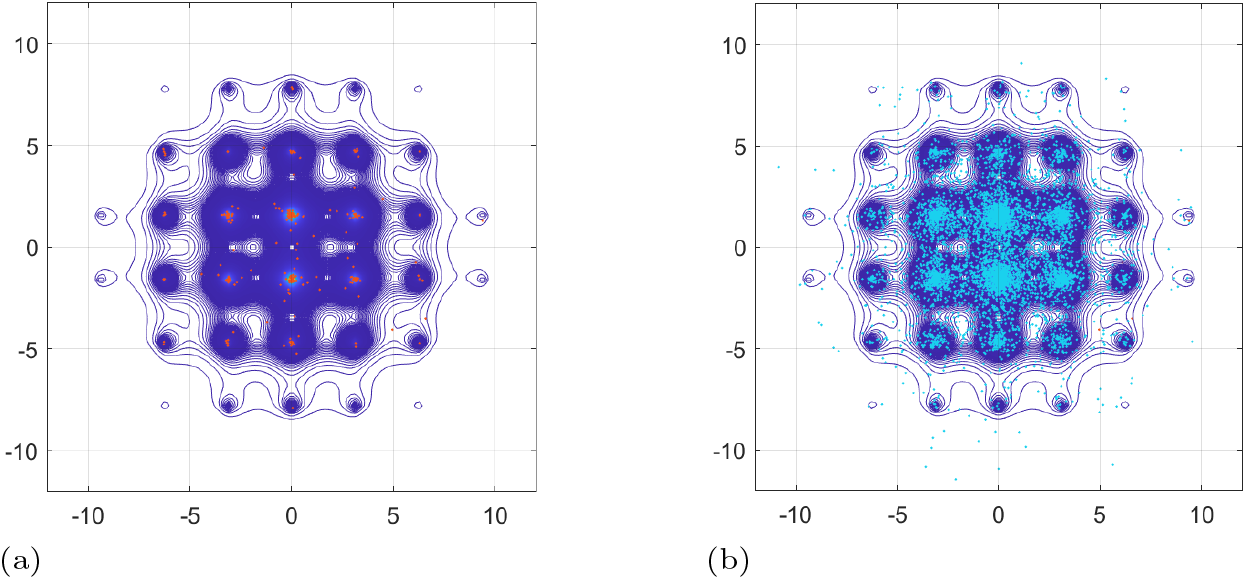
**Exp. 3**. Qualitative comparison of the numerical solution of the PDE (34) against the predictions of the SDE schemes (37a) and (37b). (a) Isolines of the final time solution *u* of the PDE (34), cf. Figure 4, superimposed with 10^4^ (final time) realisations of the SDE scheme (37a). (b) Isolines of the final time solution *u* of the PDE (34), cf. Figure 4, superimposed with 10^4^ (final time) realisations of the SDE scheme (37b). We note that the cells described by the SDE scheme (37a) are concentrated, at the final time, almost exclusively in the “valleys” of the (fixed) environment *v*, cf. Figure 4, much more than cells described by the corrected SDE scheme (37b), and much more than the final time solution *u* of the PDE (34). The results confirm that the corrected scheme (37b) offers a much better approximation of the PDE (34) than the more more intuitive scheme (37a).

**Fig. 7.**
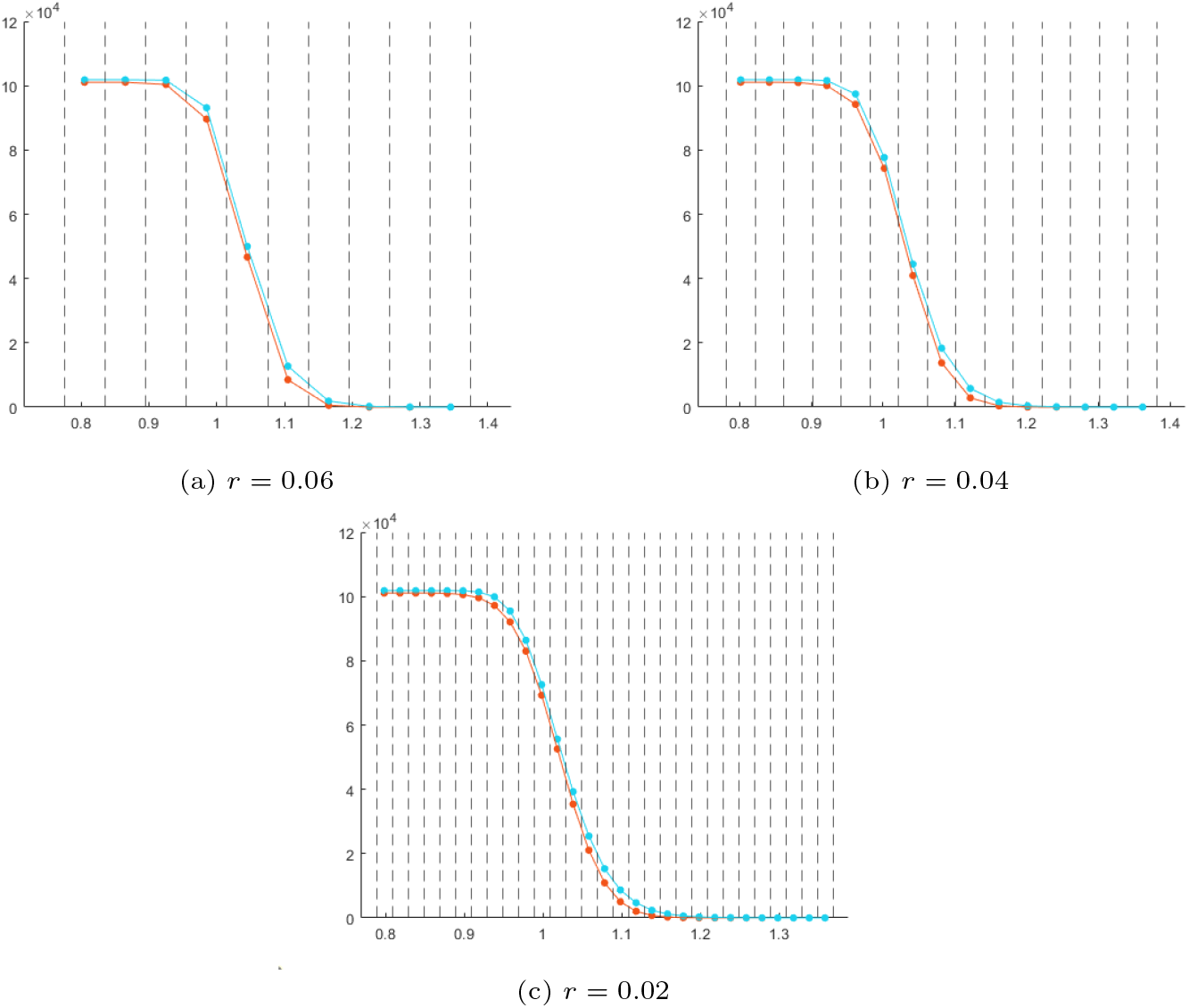
**Exp. 2**. Graphical comparison of uniform binning of the final positions of the migration tracks traversed the cells following the SDE schemes (25) and (26) respectively. For both schemes, we considered initial position at (1, 1) and a total travelling time *T* = 1. (a) Uniform binning of the final positions of migration tracks of 10^5^ cells for the SDE schemes (25) and (25) with bin size *r* = 0.06. (b) Uniform binning of the final positions of migration tracks of 10^5^ cells for the SDE schemes (25) and (25) with bin size *r* = 0.04. (c) Uniform binning of the final positions of migration tracks of 10^5^ cells for the SDE schemes (25) and (25) with bin size *r* = 0.02.

Similarly to Experiment 2, we quantify the difference between the two schemes by measuring the corresponding average distances traversed by the cells from their initial position. In more detail, we perform *K* = 10^4^ realisations for each scheme, calculate the average distances *d*_1_ and *d*_2_ using the formula (27), and their relative difference via:

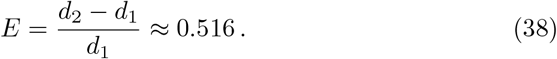

To further quantify the difference of the two schemes, we perform a uniform binning and calculate the relative error of the *J*_2_ -criterion for (37a) and (37b) for *k*–different values of the size *r* of the bins as in (32). We noticed that, for smaller values *r* we have that again 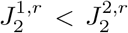. We choose *r* = 0.01 + *kh*, where *h* = 0.001, for *k* = 1, …, 140, and calculate the average value of the *k*–different values of the signed relative error *ε*^*r*^ which is

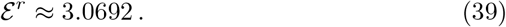

This confirms, yet again, that (37b) brings an additional migratory bias to the movement of solitary cancer cells to the entire plain.

## 5 Discussion

In this paper we have investigated the long-standing question of bridging the scales in problems of multiscale modelling and simulations. As we are motivated by the study of cancer growth and invasion models, we can rephrase this question as follows: how does one identify the correct SDEs (2) that dictate the migratory pattern of solitary cancer cells or small clusters of cancer cell, when the macroscopic evolution of the cancer cell colony follows the advection-diffusion PDE (1)?

We have exhibited in this paper, that the answer to this question is not trivial. The usual heuristic understanding that the advection and diffusion terms of the PDE (1) are responsible for the biased and random motion of the cancer cells, respectively, is not precise. Were this correct, the drift term in the SDE schemes that describe the migration of the cells in the solitary cell regime would have been solely dependent on the advection term **A** of the PDE (1), as e.g. is the case in the numerical scheme (5). We have seen with the derivations of the SDE schemes in Section 3 and with the numerical investigations in Section 4 that this is not the case. On the contrary, we have shown with (3) that the drift term of the correct SDE schemes should account for the advection **A** as well as the divergence of the diffusion **D** of the PDE (1) in a very specific way.

When comparing the corrected scheme (3) with the more heuristically expected one (5), we see that their difference depends solely on the divergence of the diffusion **D**. This clearly indicates that the two schemes would be/are identical in the case of space independent diffusion **D**. If though, both the advection and diffusion terms depend on the spatial variable—as is typically the case in cancer invasion models—then, identifying the inconsistency of the SDE schemes with the underlying PDE is not trivial and quite often is not readily apparent due to the inherent stochasticity.

From a biological/oncological perspective this is important in determining accurately how far individual cancer cells can penetrate the local tissue, and so from a clinical point of view the model results have much predictive potential. It is known that resection margins (i.e. how much tissue is removed surgically) and the pattern of cancer invasion are predictors both of local recurrence and of survival in patients who undergo surgery [21], [22], [23], [24], [25], [26]. With further refinement, accurate parameterisation and testing, using the results from the model would enable quantitative estimates to be made of the likely extent of local spread by an invasive cancer. This would then enable a surgical oncologist to tailor the radicality of surgical excision for a given individual situation. Further, more accurate estimation of metastatic spread (with the associated implications for adjuvant systemic therapy) will also be possible.

Future work in further developing the insights gained from the current modelling will focus on parameterising the model more accurately in order to make quantitative comparison with any available data. A promising avenue of development in this regard, given the difficulty in obtaining *in vivo* data, is to use *in vitro* data from organotypic invasion assays cf. [27], [28], [29].

## Supplementary Material/Appendix

### A Numerical solution of Advection-Reaction-Diffusion PDEs

We present here the main components of the numerical method we use to solve the generic *Advection-Reaction-Diffusion* (ARD) system of the form

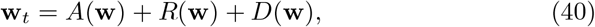

where **w** = **w**(*t*, **x**), *t* ≥ 0, **x** ∈ Ω (domain), represents the solution vector, and *A, R*, and *D* the advection, reaction, and diffusion operators respectively.

We denote by **w**_*h*_ (*t*) the corresponding (semi-)discrete numerical approximation—indexed here by the maximal spatial grid diameter *h*—that satisfies the system of ODEs

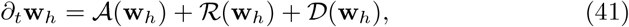

where the numerical operators 𝒜,ℜ,𝒟 and are *discrete approximations* of the operators *A, R*, and *D* in (40) respectively.

We use a second order *Implicit-Explicit Runge-Kutta* (IMEX-RK) *Finite Volume* (FV) numerical method that was previously developed in [17, 18] where we refer for more details, see also [30]. This method is based on a *splitting* in explicit and implicit terms in the form

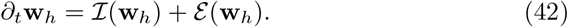

The actual splitting depends on the particular problem in hand but in a typical case, the advection terms 𝒜 are treated explicitly in time, the diffusion terms 𝒟 implicitly, and the reaction terms ℜ partly explicit and partly implicit.

After splitting, we employ a diagonally implicit RK method for the implicit part, and an explicit RK for the explicit part. They are combined in the following scheme

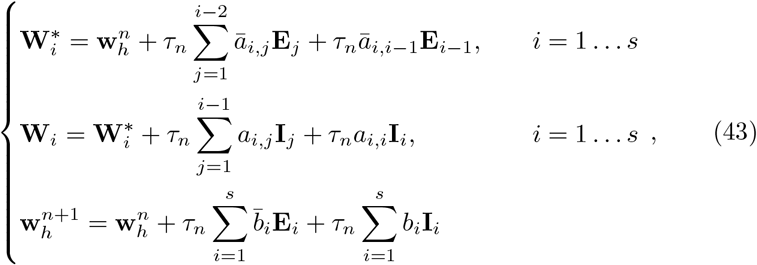

where *s* = 4 are the stages of the IMEX method, **E**_*i*_ = ***ε***(**W**_*i*_), *I*_*i*_= ℐ (**W**_*i*_), *i* = 1 … *s*,{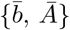, *Ā*,{*b, A*} are respectively the coefficients for the explicit and the implicit part of the scheme, given by the Butcher Tableau in Table 2, cf. [31].

We solve the linear systems in (43) using the *iterative biconjugate gradient stabilised Krylov subspace* method, for which we refer to [32, 33].

**Table 2.**
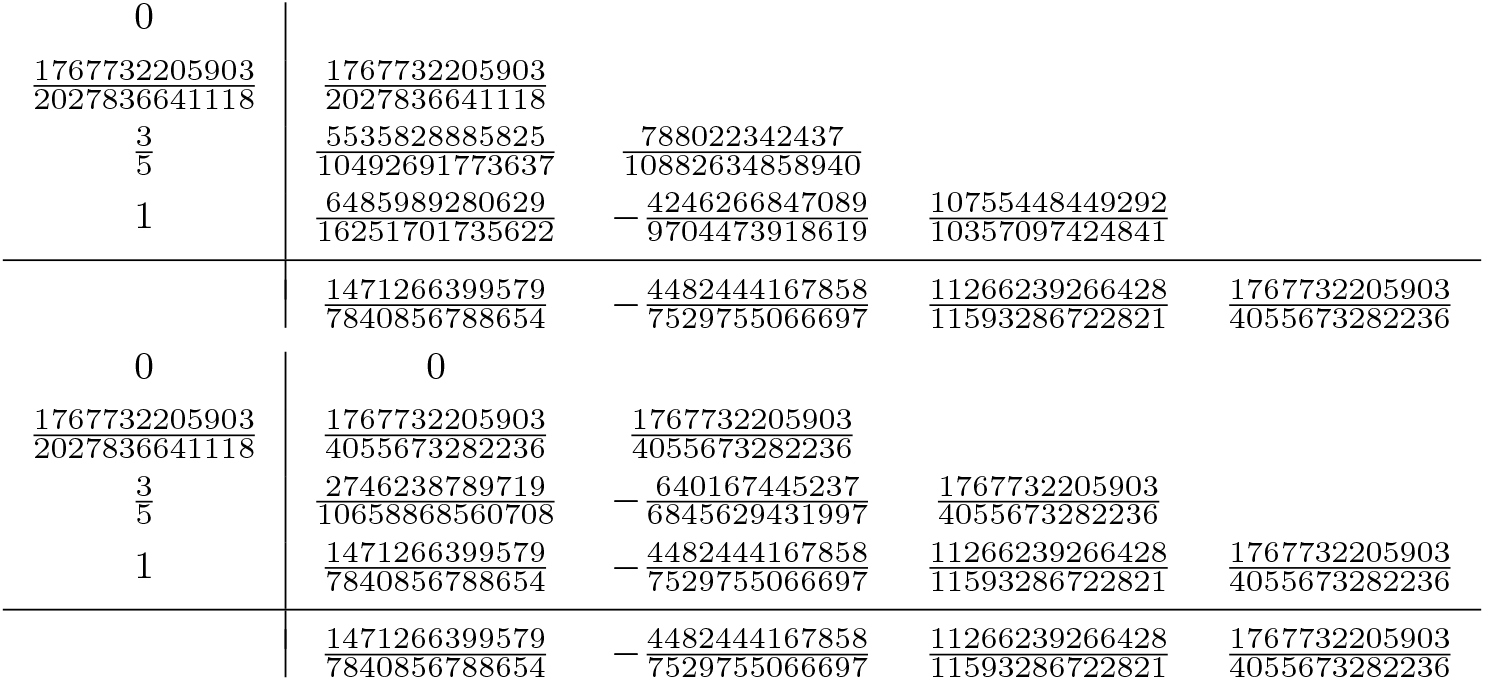
Butcher tableaux for the explicit (upper) and the implicit (lower) parts of the third order IMEX scheme (43), see also [31].

## B Uniform Binning

In this section we present some graphic results of the uniform binning performed for Experiment 2 in Section 4. We illustrate the binning of the final positions of 10^5^ realisations of both schemes (25) and (26). At the centre of each bin we place the value *U*_*i*_ calculated in (29) for both schemes. We perform this sequence with different sizes *r* of the bins. In addition we present in Table 3 the *J*_2_ values of the both schemes (25) and (26) for *r* = {0.02, 0.04, 0.06}.

**Table 3.**
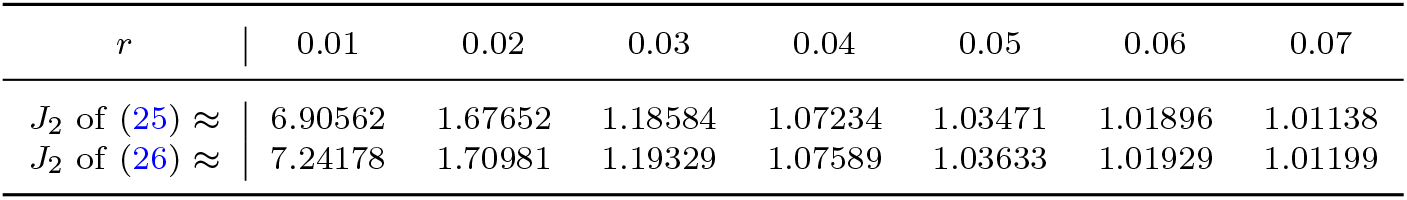
Bin size *r* and *J*_2_ values in **Exp**. 2.

| $r$ | 0.01 | 0.02 | 0.03 | 0.04 | 0.05 | 0.06 | 0.07 |
| --- | --- | --- | --- | --- | --- | --- | --- |
| $J_2$ of (25) ≈ | 6.90562 | 1.67652 | 1.18584 | 1.07234 | 1.03471 | 1.01896 | 1.01138 |
| $J_2$ of (26) ≈ | 7.24178 | 1.70981 | 1.19329 | 1.07589 | 1.03633 | 1.01929 | 1.01199 |

We also worked in a similar fashion for the schemes (37a) and (37b) in the Experiment 3.

